# Bread feeding is a robust and more physiological enteropathogen administration method compared to oral gavage

**DOI:** 10.1101/808832

**Authors:** Anne Derbise, Hebert Echenique-Rivera, Marta Garcia-Lopez, Rémi Beau, Myriam Mattei, Petra Dersch, Javier Pizarro-Cerdá

## Abstract

Oral administration is a preferred model for studying infection by bacterial enteropathogens such as *Yersinia*. In the mouse model, the most frequent method for oral infection consists of oral gavage with a feeding needle directly introduced in the animal stomach via the esophagus. In this study, we compared needle gavage to bread feeding as an alternative mode of bacterial administration. Using a bioluminescence-expressing strain of *Yersinia pseudotuberculosis*, we detected very early upon needle gavage a bioluminescent signal in the neck area together with a signal in the abdominal region, highlighting the presence of two independent sites of bacterial colonization and multiplication. Bacteria were often detected in the esophagus and trachea, as well as in the lymph nodes draining the salivary glands, suggesting that lesions made during needle introduction into the animal oral cavity lead to rapid bacterial draining to proximal lymph nodes. We then tested an alternative mode of bacterial administration using small pieces of white bread containing bacteria. Upon bread feeding infection, mice exhibited a stronger bioluminescent signal in the abdominal region as compared to needle gavage, and no signal was detected in the neck area. Moreover, *Y. pseudotuberculosis* incorporated in the bread is less susceptible to the acidic environment of the stomach and is therefore more efficient in causing intestinal infections. Based on our observations, bread feeding constitutes a natural and more efficient administration method which does not require specialized skills, is less traumatic for the animal, and results in diseases that more closely mimic food-borne intestinal infection.

## Introduction

Animal infection models represent major research tools to understand human disease, as they allow to investigate in a relevant physiological environment the very complex interactions that take place between pathogens and hosts during organ/tissue colonization or whole-body dissemination (1, 2). For the study of bacterial enteropathogens, oral administration is a preferred infection route to assess intestinal colonization and pathogenesis in mammalian hosts. The most often used methodology for oral infection in laboratories using mice as models consists of oral gavage with a feeding needle introduced in the stomach via the esophagus. However, several laboratories have reported, using bioluminescence-expressing pathogens, colonization in sites distant from the abdominal region after orogastric infection, suggesting abrasions in the laryngopharynx region (3-6). Although, the impact of such accidental colonization on the overall infectious process is not known, we cannot exclude the possibility of bacterial dissemination in the bloodstream independently to the intestinal infection. In order to avoid this potential problem, alternative modes of pathogen administration have been described, such as bread feeding (7, 8) or drinking water delivery (9, 10).

Enteropathogenic *Yersinia* are the third bacterial cause of human gastrointestinal infections in Europe (11). Although much less often isolated than *Y. enterocolitica, Y. pseudotuberculosis* is responsible for acute gastroenteritis and mesenteric lymphadenitis in a wide variety of animals including rodents, domestics animals, non-human primates and humans (12). After oral ingestion, *Y. pseudotuberculosis* localizes to the ileum and proximal colon and can pass/cross the intestinal barrier, invading its host through gut-associated lymphoid tissues and Peyer’s patches (13). In humans, dissemination to deeper tissues and the bloodstream are frequent in patients with underlying disease conditions (such as diabetes mellitus, liver cirrhosis, and hemochromatosis) and can lead to septicemia as a severe outcome of the infection (14) (15).

While performing mice oral administration of *Y. pseudotuberculosis* with a feeding needle, we regularly noticed that some animals exhibit spleen infections without displaying the presence of bacteria in Peyer’s patches (PPs) or mesenteric lymph nodes (MLNs), suggesting the passage of bacteria in the blood stream independently of the colonization of the gut-associated lymphoid tissues (data not shown). Since it has been reported that needle gavage can cause lesions in the oropharyngeal region (3), we decided to re-examine this mode of administration and compare it to an alternative administration method (bread feeding) with a fully virulent *Y. pseudotuberculosis* strain expressing bioluminescence in order to follow bacterial dissemination over time in the whole animal body. Our results clearly illustrate that needle feeding promotes *Y. pseudotuberculosis* colonization not only of the intestinal tract but also the neck region with a tropism for salivary glands lymph nodes. On the contrary, bread feeding induces a much more robust intestinal tract infection that is never associated to neck region infections. Moreover, bread feeding allows to protect bacteria from the acidic environment of the stomach. We conclude therefore that bread feeding is a better oral administration route to investigate the pathophysiology of bacterial enteropathogen infection.

## Results

### Construction of a constitutively bioluminescent *Y. pseudotuberculosis* strain

The IP32953 *Y. pseudotuberculosis* strain, isolated from the stools of a human patient, was chosen for this study due to its capacity to express two major virulence factors, the type 3 secretion system (T3SS) and the Yersiniabactin (Ybt) iron uptake machinery. This strain has been previously shown to be fully virulent in laboratory mice (16) (17).

To perform a comparative analysis of bacterial dissemination *in vivo* after oral infection, the strain IP32953 was genetically engineered to constitutively express bioluminescence. In order to correlate bioluminescence to bacterial numbers during animal infection, the *luxCDABE* operon (placed under the control of the constitutive *rplN* promoter) was stably introduced into the *Y. pseudotuberculosis* chromosome using the mini-Tn7-transposon technology (18, 19). The resulting strain, IP32953-lux, harbors the construct *mini-Tn7-Km-P*_*rplN*_*-luxCDABE* between the two housekeeping genes *glmS* and *pstS* (**Fig. S1A**)(19). Stability of the bioluminescence signal over time was determined by performing 10 subcultures of IP32953-*lux* in LB without antibiotics over 17 days. At different time points, bacterial aliquots were streaked on LB plates and individual colonies were checked for their capacity to emit bioluminescence. After 10 subcultures, 100% of the CFU expressed bioluminescence confirming the stability of the bioluminescent phenotype in the bacterial population (data not shown).

Strain IP32953-*lux* displayed similar *in vitro* growth and virulence as the parental wild type IP32953 strain (data not shown). Therefore, IP32953-*lux* was used to determine the characteristics and relevant differences between the two infection methods applied: oral gavage with a feeding needle versus bread feeding.

### *Y. pseudotuberculosis* colonizes the neck and the abdominal region of infected animals upon needle feeding

We first set up the bread feeding protocol in OF1 mice based on a previously reported methodology (7). In our study, instead of using melted butter as a vehicle to deliver bacteria and to attract mice to bread, mice were first trained to feed on bread only (see Materials & Methods) and then were infected with bread supplemented with a bacterial suspension in PBS. A pilot experiment using this new method allowed us to validate that within less than 5 min mice consumed bread supplemented with 8E7 IP32953-*lux* CFUs, subsequently exhibiting bioluminescence in the PPs and MLNs, confirming therefore *Y. pseudotuberculosis* colonization and dissemination in the intestinal tract as expected for an enteropathogen **(Fig. S1B).**

We then proceed to compare needle versus bread feeding as oral administration routes. 24 h after oral administration of 3.5E8 IP32953-*lux* CFUs, mice were monitored for bioluminescence and regions of interest (ROI) were identified for bioluminescence imaging (BLI) measurements. Upon needle gavage, up to 80% of the mice displayed a BLI signal in two distinct regions of the body, one in the expected abdominal region and a second in the neck, whereas upon bread feeding none of the mice exhibited a neck signal and bioluminescence was restricted to the abdomen (**Fig. 1A**). ROI measurements in the neck region of needle-infected mice indicate that the level of BLI is similar to the one coming from the abdominal region, suggesting that bacteria colonize, disseminate and multiply in the neck as efficiently as in the abdomen (**Fig. 1B**). This comparative analysis indicates that oral administration of *Y. pseudotuberculosis* using needle feeding results in infection of the neck region, a phenotype that is not observed upon bread feeding.

**Fig. 1.**
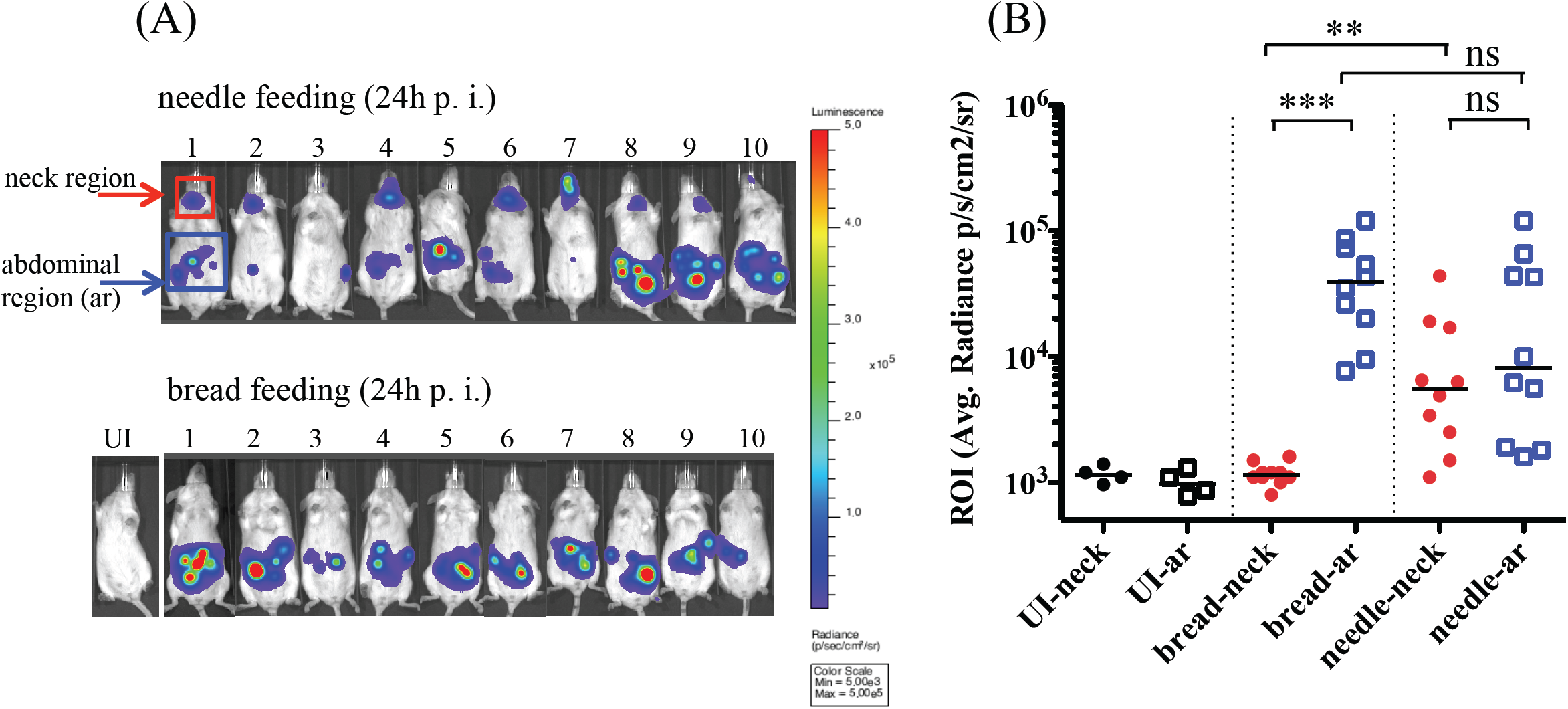
Comparative bioluminescence imaging of mice infected by either needle or bread feeding. OF1 mice were infected with 3.5E8 CFUs of *Y. pseudotuberculosis* bioluminescent strain (IP32953-*lux*) using a 20G x 1,5” feeding needle or a piece of bread. (A) At 24 hours (h) post infection mice were imaged using an IVIS Spectrum imaging system with an acquisition time of 2 minutes and small binning. Uninfected mice (UI) were used to set the light emission background. Regions of interest (ROI) were drawn in the neck region (red frame) and the abdominal region (ar, blue frame) using the Living Image 4.5 software. (B) ROI average bioluminescence (photon/sec/cm2/sr) was calculated for each individual mice and data were analyzed using Prism 5.0 software for T test non-parametric Mann Whitney, p>0,05(ns), p<0,0015(**), p<0,0002(***). Median of the values are indicated by an horizontal bar. Bioluminescence signal is detected in the neck of 80% of the needle infected animals with an average radiance not statistically different than the signal from the abdominal region, whereas none of the mice infected with bread presented a signal in the neck. Although the median of the abdominal region is higher when mice are infected with bread, the overall signal is not statistically different due to the variability between mice.

### Upon needle feeding, *Y. pseudotuberculosis* multiplies in the intersection of the oesophagus and trachea and disseminates to the draining lymph nodes of the submandibular area

To identify which specific neck region is colonized by *Y. pseudotuberculosis*, mice were first orally gavaged with 4E8 IP32953-*lux* CFUs; then at 24 and 48 h post infection, mice exhibiting a BLI signal in the neck were euthanized and dissected in the cervical ventral region (see Materials & Methods). After skin removal, all the mice exhibited a BLI signal coming from the cervical soft tissue composed of salivary glands, lymph nodes (LNs) and adipose tissues (**Fig. 2**). Among 10 dissected mice, nine exhibited bioluminescent signals in LNs from the salivary glands region (**Fig. 2A, 2C_1_, 2D_1_**). Although there was no preferential right or left LN colonization, we often noticed that after removal of a first LN producing high amounts of photons, it was possible to identify secondary LNs producing less light, indicating variable levels of bacterial colonization among LNs. Each time a BLI positive LN was isolated, we verified its bacterial content by homogenization and CFU counting on agar plates (**Fig. 2C_1_, 2D_1_**). In addition to LNs, we noticed in 70% of the mice a strong BLI signal in the esophagus and/or trachea (**Fig. 2B, 2C**). Dissection of the esophagus and trachea sections associated with BLI allowed us to localize bacterial colonization at the junction of these two structures (**Fig. 2C_2_**), suggesting a probable deposition of bacteria in the tissue consecutive to the introduction of the needle in the esophagus. Finally, half of the mice emitted a BLI signal in the oral cavity corresponding most of the time to the skin associated to the lip (**Fig. 2D, 2D_2_**). From these results, we speculate that the use of a feeding needle entails a risk of creating lesions in the skin of the mouth as well as in the tissue at the junction of the esophagus and trachea, from which *Y. pseudotuberculosis* can disseminate to and multiply in the draining LNs located in the salivary glands region.

**Fig. 2.**
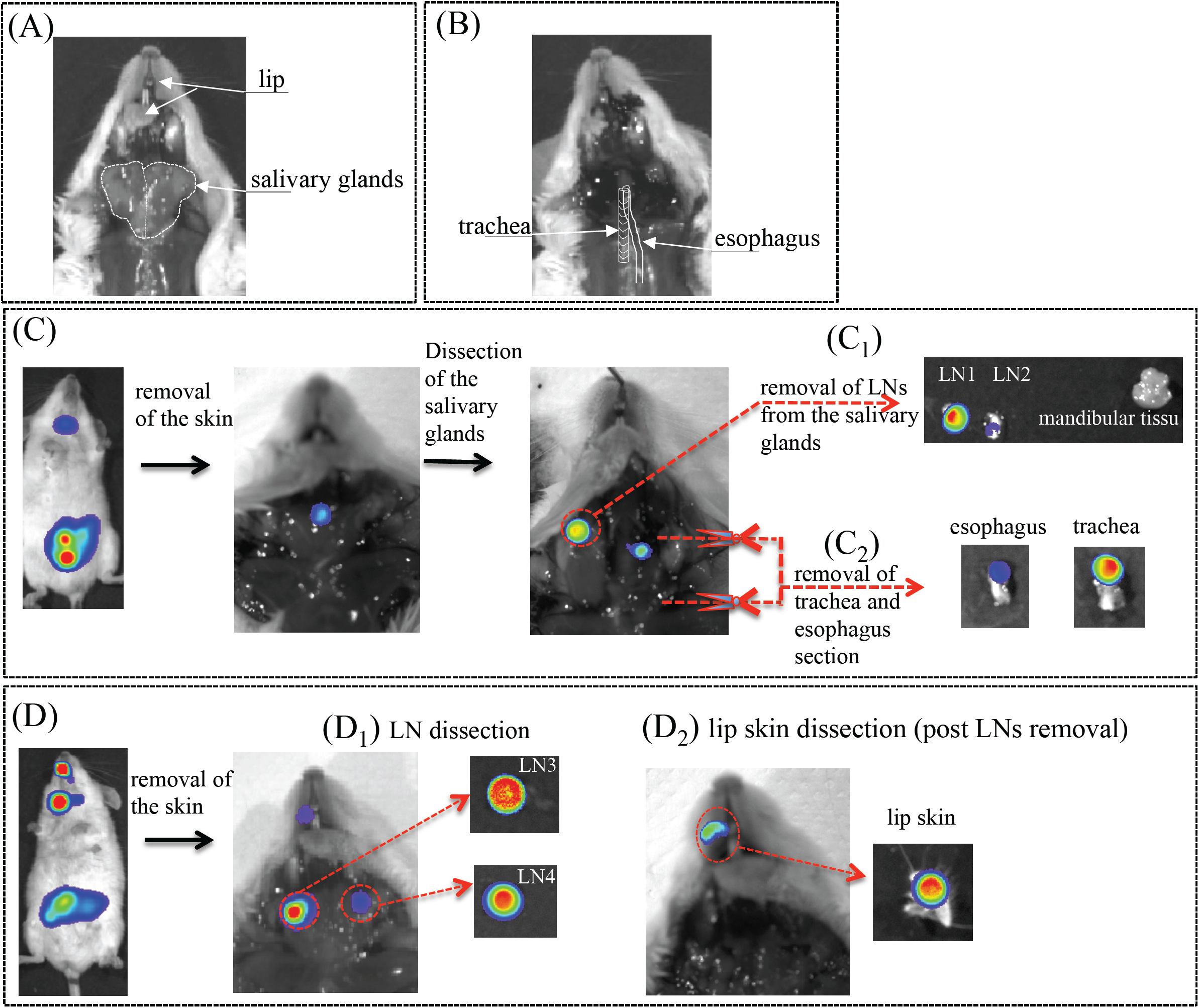
Mouse regional ventral cervical anatomy and analysis of the site of bacterial colonization after needle feeding. (A) salivary glands outlined in white doted lines are bilateral located in the cervical neck, (B) after the salivary glands removal, trachea and underneath esophagus are accessible as schematically indicated. (C) and (D) Mice exhibiting BLI signal in the neck 24 h or 48 h post-infection (needle feeding with 4E8 CFU of IP32953-*lux*) were euthanized using CO_2_ and a dissection procedure associated with BLI acquisition was performed step by step. Two representative mice are shown. After a first step of skin removal, the salivary glands region was dissected and lymph nodes (LN1-4) found in this region were collected. Then the esophagus and trachea section were collected as indicated by the schematized red scissors. Regions of interest (ROI) were drawn and average bioluminescence (photon/sec/cm2/sr) was calculated for each dissected organs. Organs exhibiting BLI signal were homogenized in PBS and enumerated for bacterial count (CFU). Panel C_1_: lymph nodes : LN1(ROI=2E5; CFU=2E4), LN2 (ROI=5.4E4; CFU=4E3). Panel C_2_: esophagus (ROI=2E6; CFU=4E5) and trachea (ROI=6E6; CFU=6E5). Panel D_1_ : lymph nodes LN3 (ROI=2.5E7; CFU=4E6) and LN4 (ROI=1E6; CFU=4E5). Panel D_2_: skin from the mouth (ROI=9.3E5; CFU=not done). Among 10 dissected mice, nine exhibited bioluminescent signals in lymph nodes from the salivary glands region (panel C_1_ and D_1_), 7 in the esophagus and/or trachea (panel C_2_) and five close to the mouth/lip region (panel D_2_).

Although we do not know the impact of the *Y. pseudotuberculosis* neck colonization in the overall infectious process, we cannot exclude the possibility of bacterial dissemination in the blood stream occurring independently to the intestinal colonization and translocation at the gastro-intestinal barrier.

### Bread feeding protects *Y. pseudotuberculosis* from the acidic gastric environment and promotes efficient intestinal colonization

We next compared the kinetics of bacterial dissemination upon bread feeding versus needle feeding. Mice were orally infected with 3.5E8 IP32953-*lux* CFUs and imaged at 0.5, 6, 24, 48 and 72 h post infection (**Fig. 3**). At each time point, photon emission was measured from the abdominal region and neck region. As indicated in **Fig. 3A** and **3B**, upon bread feeding the BLI signal in the abdominal region is detected as early as 30 min post infection, then decreases to an almost undetectable signal at 6 h p.i., and in a second phase increases continuously. Upon needle feeding the BLI signal is barely detectable before 24 h p.i.: after that time point, the BLI signal is detected in the neck **(Fig 3A, 3D)** and the abdominal region **(Fig. 3A, 3B)** where it continuously increases over time. It is noteworthy that the neck region signal was never detected in bread-infected animals **(Fig. 3D)** even at later time points (data not shown). Statistical analysis indicates a significantly higher light emission signal in the abdominal region at early (0.5 h) and later time points (72 h) when using bread compared to needle **(Fig. 3B).** This observation correlates with the lower amount of *Y. pseudotuberculosis* found in the feces six hours post needle feeding as shown by CFU counting (**Fig. 3C**), suggesting a lower number of bacteria surviving the passage through the stomach.

**Fig. 3.**
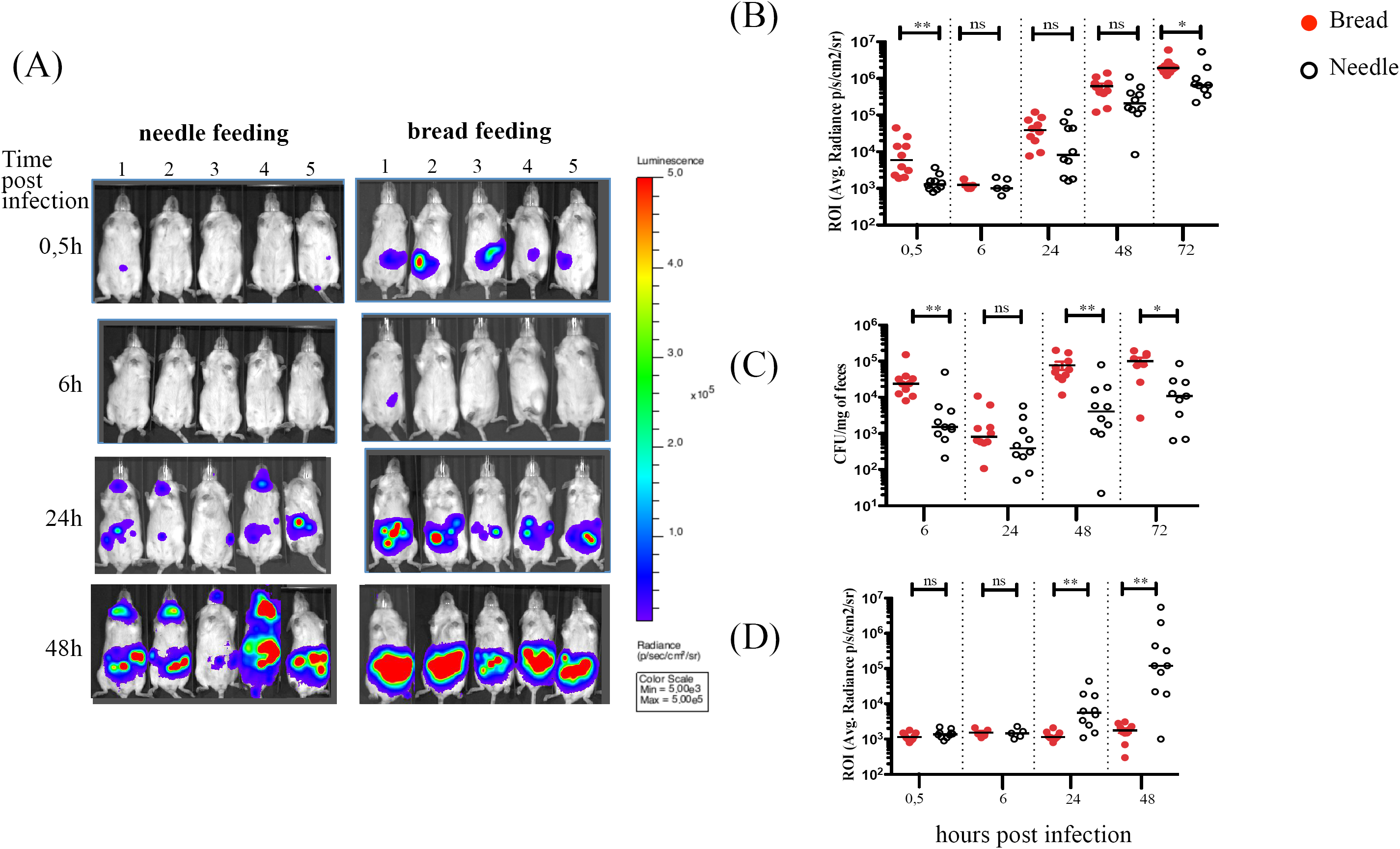
Comparative analysis of bacterial survival upon bread versus needle feeding. OF1 mice were infected with 3,5E8 IP32953-*lux Y. pseudotuberculosis* CFUs using bread or feeding needle and at 0.5, 6, 24, 48 and 72 h post infection mice were imaged using an IVIS Spectrum imaging system. (A) Monitoring of 5 representative mice from 0.5 to 48 h post infection using needle and bread feeding, same color scale (min=5E3 max=5E5) with settings 2 min time of exposure and small binning. (B) ROI were drawn in the abdominal region and average bioluminescence (photon/sec/cm2/sr) was calculated for each mouse at 0.5, 6, 24, 48 and 72 h post infection using bread (red circle) or needle (open circle). (C) Enumeration of *Y. pseudotuberculosis* IP32953-*lux* in feces at 0.5, 6, 24, 48 and 72 h post infection using bread (red circle) or needle (open circle) feeding. (D) ROI were drawn in the neck region and average bioluminescence (photon/sec/cm2/sr) was calculated for each mouse at 0.5, 6, 24, 48 and 72 h post infection using bread (red circle) or needle (open circle) feeding. Median of the values are indicated by a horizontal bar. Data were analyzed using Prism 5 software for T test non-parametric Mann Whitney, p<0,002 (**); p=0,014(*); ns: not significative p>0,05. The increased BLI signal in the abdominal region and bacterial load in the feces when mice are infected by bread feeding indicates a more efficient delivery of bacteria and colonization of the intestinal tract compared to the needle feeding protocol. BLI signal increases over time in the neck of mice infected with the needle while no signal was detected in the neck of the animals infected with bread.

When needle feeding is used, bacteria are delivered directly to the stomach, a compartment known to have an acidic pH aggressive for microorganisms. To evaluate whether the lower amount of light observed in the abdominal region of needle-infected mice was due to bacterial killing by the acidity of the stomach, we infected animals using the needle with either bacteria resuspended in PBS only, or in PBS buffered with CaCO_3_ (20). As shown in **Fig. 4A** and **4B**, buffering the bacterial inoculating medium significantly increases the BLI signal in the abdominal region of infected mice as well as the amount of bacterial CFUs recovered in the feces (**Fig. 4C**). Our results therefore indicate that bacterial bread delivery protects *Y. pseudotuberculosis* from the acidic gastric environment and promotes efficient intestinal colonization.

**Fig. 4.**
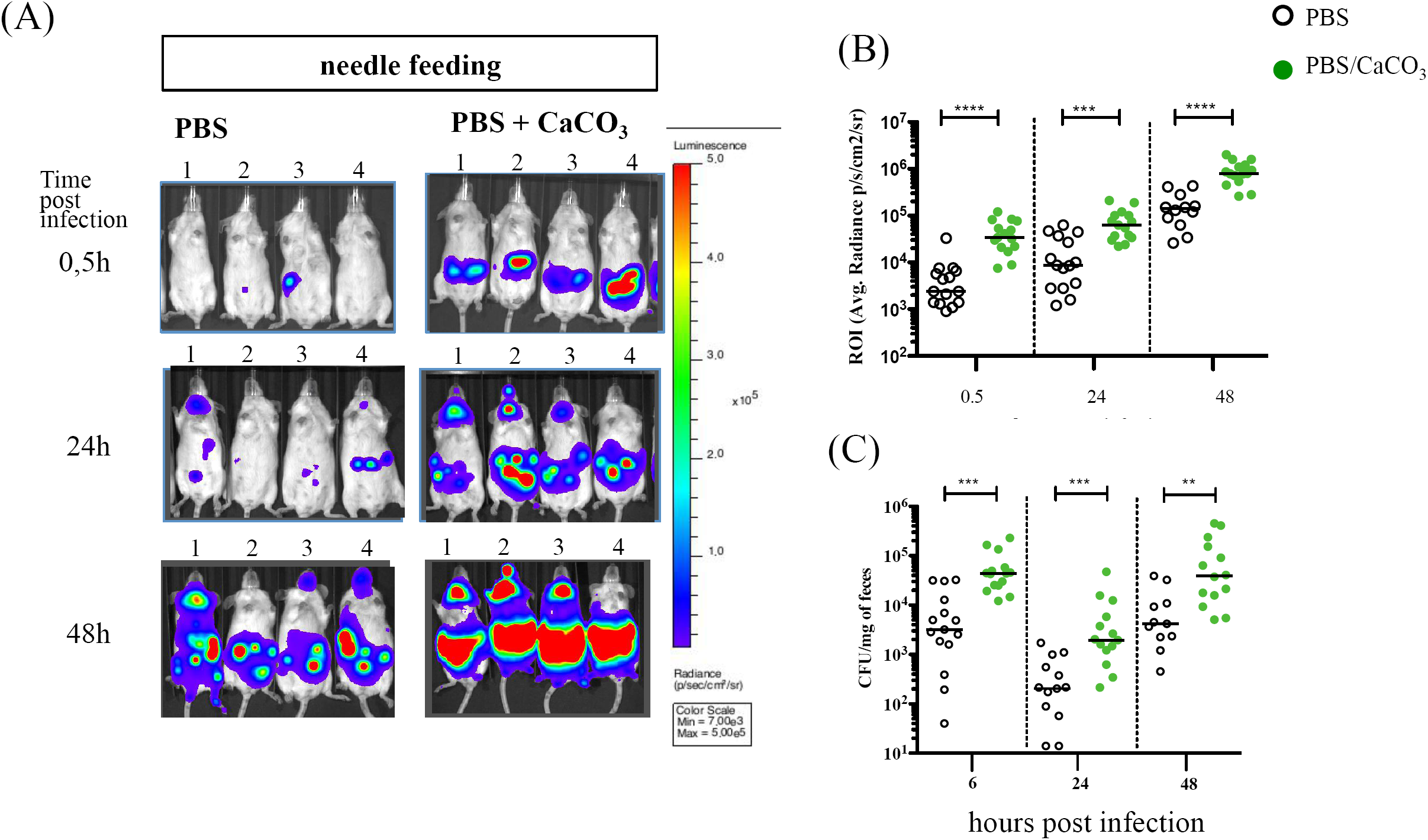
Protective effect of CaCO_3_ when bacteria are administered directly in the stomach by needle feeding. OF1 mice were infected with 4 to 5E8 *Y. pseudotuberculosis* IP32953-*lux* CFUs using two needle feeding conditions: bacterial suspension in PBS (open circle) or in PBS supplemented with CaCO_3_ (30 mg/ml) (green circle). At 0.5, 24 and 48 h post infection mice were imaged using an IVIS Spectrum imaging system. (A) A representative panel of the animals for the two conditions are shown using same color scale (min=7E3 max=5E5) with settings 2min time of exposure and small binning. (B) Regions of interest (ROI) were drawn in the abdominal region and average bioluminescence (photon/sec/cm2/sr) was calculated for each mouse. (C) Enumeration of *Y. pseudotuberculosis* IP32953-*lux* in feces at 6, 24, and 48 h post infection. Median of the values are indicated by a horizontal bar. Data were analyzed using Prism 5 software for T test non-parametric Mann Whitney, p<0,0001 (****), p>=0,0001(***); p=0,0014(**). Addition of CaCO_3_ to the bacterial suspension protects *Y. pseudotuberculosis* when administered via a feeding needle directly in the stomach.

### Mice are more susceptible to *Y. pseudotuberculosis* infection when administered via bread feeding

The better survival of *Y. pseudotuberculosis* when associated with CaCO_3_ prior to needle feeding led us to evaluate whether it would change the overall lethal dose 50 (LD_50_) of strain IP32953-*lux* when administered with bread. Thus, mice were infected using three conditions (needle with or without supplementation of CaCO_3,_ and bread) with four different *Y. pseudotuberculosis* IP32953-*lux* concentrations (2.5E5, 2.5E6, 2.5E7, and 2.5E8 CFU). Animals were monitored over time for BLI imaging (**Fig. 5 and 6**), body weight loss (**Fig. 5B, 6B**), signs of disease, and lethality to allow measurements of LD_50_ (**Fig. 7**).

**Fig. 5.**
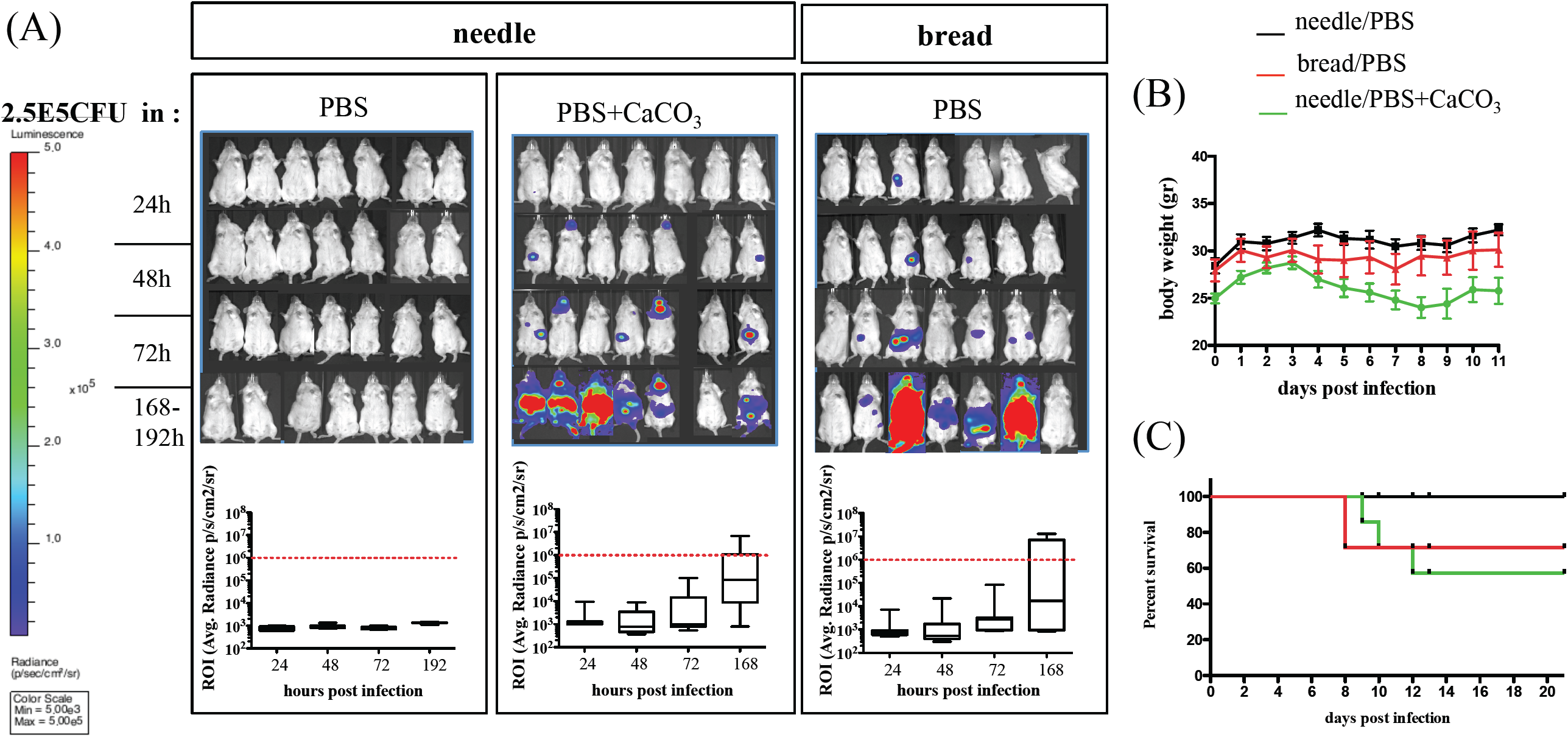
Comparative analysis of the course of infection using the three protocols of administration with 2.5E5 CFU of *Y. pseudotuberculosis*. OF1 mice were orally infected with 2.5E5 (A, B, C) *Y. pseudotuberculosis* strain IP32953-*lux* CFUs and monitored overtime for BLI signal, weight loss and mortality after needle feeding of bacteria resuspended in PBS (black line), in PBS supplemented with CaCO_3_ (green line), or after bread feeding of bacteria resuspended in PBS only (red line). Quantification of the bioluminescent signal from the abdominal region is shown below the mice pictures in each condition at each time point (24, 48, 72 and 168-192 h p.i.). Red dotted lines indicate the average radiance of ROI measurements above which animals are in terminal illness. Red crosses positioned on the X axis indicate dead mice. Comparative weight loss (B) and survival of mice (C) for 21 days using the three protocols of administration. Bread feeding allows a more efficient infection of the abdominal region by *Y. pseudotuberculosis* without neck colonization compared to the needle feeding.

**Fig. 6.**
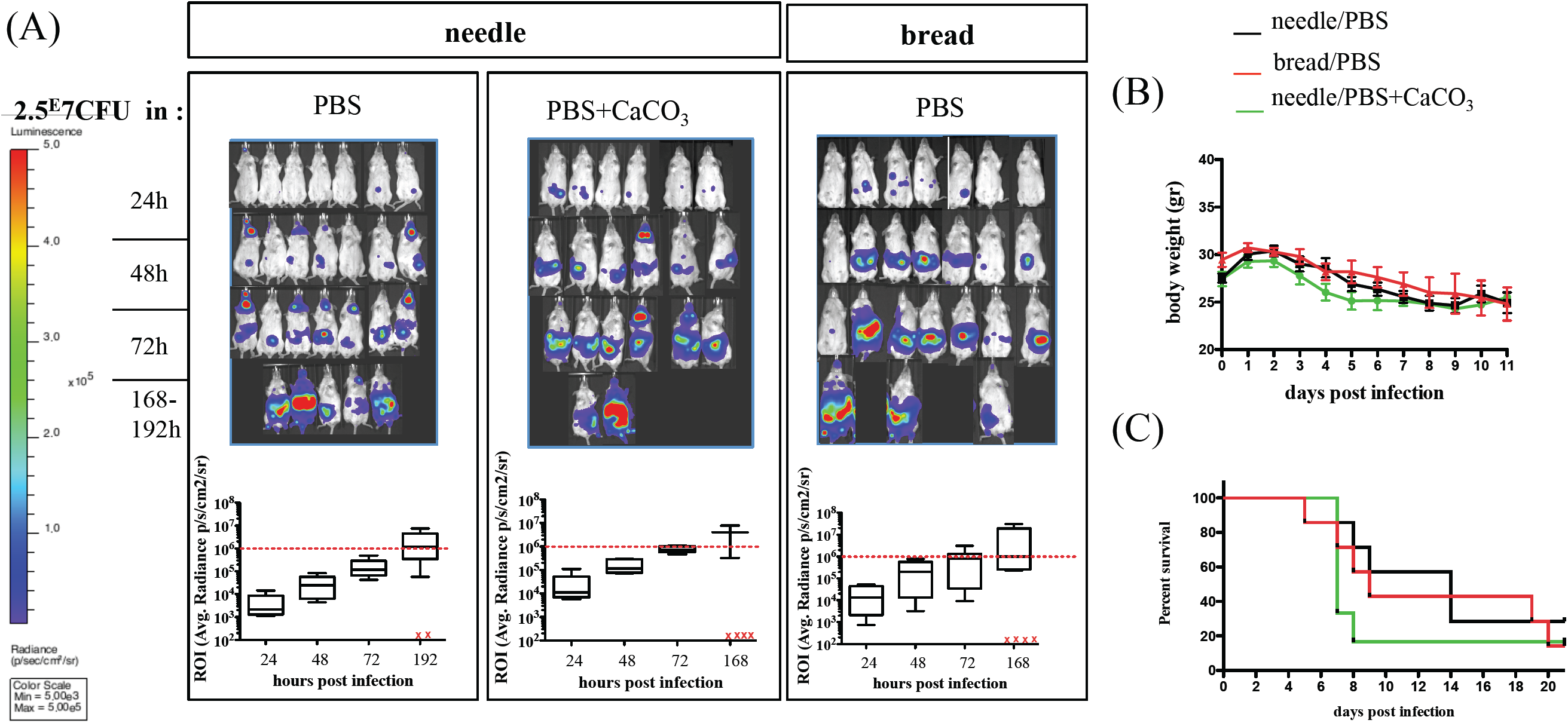
Comparative analysis of the course of infection using the three protocols of administration with 2.5E7 CFU of *Y. pseudotuberculosis*. OF1 mice were orally infected with 2.5E7 (A, B, C) *Y. pseudotuberculosis* strain IP32953-*lux* CFUs and monitored overtime for BLI signal, weight loss and mortality after needle feeding of bacteria resuspended in PBS (black line), in PBS supplemented with CaCO_3_ (green line), or after bread feeding of bacteria resuspended in PBS only (red line). Quantification of the bioluminescent signal from the abdominal region is shown below the mice pictures in each condition at each time point (24, 48, 72 and 168-192 h p.i.). Red dotted lines indicate the average radiance of ROI measurements above which animals are in terminal illness. Red crosses positioned on the X axis indicate dead mice. Comparative weight loss (B) and survival of mice (C) for 21 days using the three protocols of administration.

**Fig. 7.**
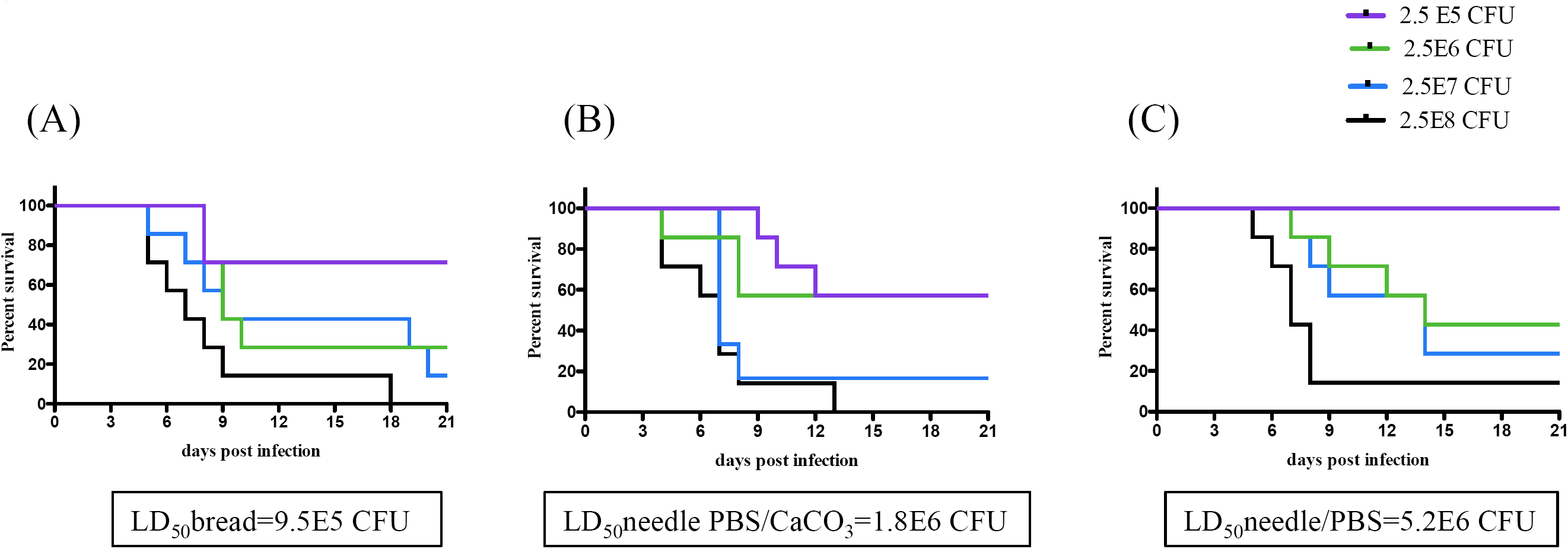
Comparative analysis of animal survival after oral infection with *Y. pseudotuberculosis*. OF1 mice survival after oral infection with serial dilutions of *Y. pseudotuberculosis* IP32953-*lux* using bread feeding (A), needle feeding supplemented with CaCO_3_ (B) or needle feeding without CaCO_3_ (C). Mice were observed daily and LD_50_ were determined according to the method of Reed and Muench (21).

The use of the needle without supplementation of CaCO_3_ gave the highest LD_50_ (5.2E6 CFU)(**Fig. 7C**). When bacteria are mixed with CaCO_3_, the LD_50_ is lower (1.8E6 CFU) suggesting a beneficial effect of CaCO_3_ when the needle feeding administration is used (**Fig. 7B**). However, the lowest calculated LD_50_ was obtained when OF1 mice were bread-infected (LD_50_=9.5E5) (**Fig. 7A**). The BLI and weight loss illustrate very well the differences of bacterial dissemination and multiplication at a low dose (2.5E5 IP32953-*lux* CFUs) between the three infection conditions (**Fig. 5A, 5B**). At this low dose, none of the mice treated with the needle (without supplementation of CaCO_3_) showed signs of disease or weight loss while the mice treated with bread or needle supplemented with CaCO_3_ showed signs of disease, weight loss and mortality (30 to 40% of them died) **(Fig. 5B, 5C).** When the same parameters are monitored after oral infection with a higher dose (2.5E7 IP32953-*lux* CFUs) no significant differences were observed between the 3 conditions of administration, as shown in **(Fig. 6A, 5B, 5C**). It is noteworthy that even at low bacterial concentrations (2.5E5 CFU), needle feeding (with or without CaCO_3_) leads to neck colonization, whereas none of the mice showed a neck BLI signal upon bread feeding (**Fig. 5A**). Our results therefore show that infection using bread feeding is more efficient for *Y. pseudotuberculosis* delivery to the intestinal tract, resulting in better colonization even at low inoculum concentrations.

## Discussion

When studying host pathogen interactions, the choice of a laboratory experimental model of infection is crucial. Besides the choice of the animal species and the pathogen to study, important parameters that have to be taken into account include the control of the dose of the infectious agent administered to each animal, the rapidity to handle the animal and the stress induced by the animal manipulation. For decades, laboratories interested in studying animal oral infection have used needle feeding to orally deliver infectious agents. Although needle feeding allows to control the delivered dose, there are drawbacks such as the requirement of a specific manipulator training, the use of anesthesia when animals are particularly agitated, or the stress induced by the handling of the animal. Importantly, in this study we show that the use of needle gavage induces damage to the oral cavity of infected animals, leading to pathogen colonization in tissues and organs distant from the intestinal tract. Although it is not known whether such artificial colonization affects the intestinal infection process, it is reasonable to question its impact on the overall host immune response.

In order to avoid this potentially unwanted response, we propose the use of a bread feeding methodology which allows a non-traumatic pathogen administration, where animal handling is minimized and therefore is less stressful for both the manipulator and the animal. We found that the habituation step to feed on bread few days prior to the infection is crucial for effective bread feeding, with no need of additional melted butter as a vehicle as proposed by others (7) (8).

Bread feeding constitutes a very good administration method since it allows a control of the dose administered and is a fast procedure where all bread is generally consumed by mice in less than 5 minutes as opposed to the drinking water method (within several hours) described by others (10).

In addition, our study shows that *Y. pseudotuberculosis* colonizes better the intestinal tract when administered with bread compared to the needle, with no need to buffer the bacterial cell suspension. The differences in LD_50_ measurement obtained in our study using the same *Y. pseudotuberculosis* virulent strain but different administration protocols encourage us to reinvestigate the infectious process by *Y. pseudotuberculosis* after bread feeding delivery. Finally, we expect that this delivery method can be extended to other studies of host pathogen interactions with different intestinal pathogens.

## Materials and Methods

### Culture conditions and bacterial strain construction

Bacteria were grown at 28°C in lysogeny broth (LB) or on LB agar (LBA) plates. Bacterial concentrations were evaluated by spectrometry at 600 nm and plating on LBA. Kanamycin (Km, 30 μg/ml), or irgasan (0.1 μg/ml) were added to the media when necessary.

The fully virulent IP32953, serotype 1b *Y. pseudotuberculosis* isolated from a human stool in France, was used as parental strain to generate IP32953-*lux*, a bioluminescence expressing strain. As previously described (19) the *Photorabdus luminescence lux CDABE* operon controlled by the *rplN* constitutive *Yersinia* promoter was introduced into IP32953 chromosome via the Tn*7* technology (18). Plasmid pUCR6K-mini-Tn*7*-*Km*^*r*^*-luxCDABE* and the transposase-encoding plasmid pTNS2 were co-electroporated to IP32953 electrocompetent cells. Bioluminescent recombinant clones were selected on LB agar plates supplemented with Km. The recombinant *Y. pseudotuberculosis* IP32953-*lux* was verified (i) for Tn7-P*rplN*-lux chromosomal insertion by PCR using primers flanking the Tn*7* insertion site, P*glmS* 5′-gctatacgtgtttgctgatcaagatg-3′, P*pstS* 5′-acgccaccggaagaaccgatacct-3′, PTn*7*L 5′-attagcttacgacgctacaccc-3′, and PTn*7*R 5′-cacagcataactggactgatttc-3′, (ii) for similar growth rate as the parental strain in LB, and (iii) for similar virulence via oral route using the needle feeding administration. Stability of the *lux* operon was verified by successive subcultures in LB without antibiotic pressure and measurements of BLI signal on CFU resuspended in 0.1 mL LB using a Xenius 96 well plate reader (SAFAS Monaco).

### Ethics statement

Animals were housed in the Institut Pasteur animal facility accredited by the French Ministry of Agriculture to perform experiments on live mice (accreditation B 75 15-01, issued on May 22, 2008), in compliance with French and European regulations on the care and protection of laboratory animals (EC Directive 86/609, French Law 2001-486 issued on June 6, 2001). The research protocol was approved by the French Ministry of Research (N° CETEA 2014-0025) and Institut Pasteur CHSCT (n°0399).

### Animal experiments

Female 7-week-old OF1 mice were purchased from Charles River France and allowed to acclimate for 1 week before infection. Prior to infection mice were fasted 16 h and had continuous access to water. All oral infections were performed on non-anesthetized animals. Serial dilution of bacterial suspension was performed in Phosphate-buffered saline (PBS without CaCl_2_/MgCl2) from cultures grown for 48 h at 28°C in LB agar plates.

For oral gavage mice were administered a 0.2 mL bacterial suspension using an animal feeding stainless steel bulbous-ended needle (0.9 mm × 38 mm, 20G × 1.5″, Cadence Science cat. no. 9921). The bulbous-ended needle was inserted over the tongue into the esophagus and stomach as previously described (20). When required, the 0.2 ml bacterial suspension used for oral gavage (feeding needle) was mixed with 0.3 ml of a 50-mg ml^−1^ suspension of CaCO_3_ in PBS without CaCl_2_/MgCl_2_ and the 0.5 ml mix was administered. Since CaCO_3_ is not soluble at this concentration the bacterial suspension was mixed to CaCO_3_ at once before each feeding needle injection.

For bread feeding, mice were first adapted to feed on bread prior to the infection. Thus, three days before infection, the food was replaced by small pieces of white bread (approximately 9 mm^2^) to allow mice to feed on bread for a 2 h period. The same bread adaptation was repeated once 24 h before infection. Then 16 h prior to the infection the food was removed, and mice were fasted with access to water. A 20 μL bacterial suspension (without supplementation of CaCO_3_) was deposited on one piece of bread, placed in an empty and clean cage where one mouse was introduced. Each mouse was visually monitored until complete bread feeding. Generally, bread feeding took from 30 seconds up to 10 minutes per mouse. After feeding animals were housed in a cage with new litter, and access to food and water *ad libitum*.

After infection animals were monitored daily for 21 days and every day the litter was renewed in order to limit accumulation of feces in the cage and avoid cross contamination between mice.

### BLI imaging and dissection

*In vivo* imaging was performed with an In Vivo Imaging System (IVIS 100, Caliper Life Sciences). Animals were anesthetized using a constant flow of 2,5% isoflurane mixed with oxygen. Images were acquired with binning 4 and an exposure time from 10 sec. to 2 min. according to the signal intensity. To quantify luminescence signal, region of interest (ROI) were drawn and measurements of the ROI are given as average radiance (photons/s/cm2/steradian). Uninfected mice were used to set the light emission background. Analysis of the cervical region was performed sequentially starting by the removal of the skin on euthanized animals and imaging, followed by removal of the most intense bioluminescent tissue and imaging again of the animal to identify lower intensity signal. When signal was still detected after removal first identified bioluminescent tissue the same sequence was repeated until no signal could be detected. Each organ and tissues were aseptically removed, placed in glass beads containing tubes 0.5 ml PBS and subjected to homogenization to determine bacterial loads. Feces were collected from live mice and were homogenized in PBS using disposable homogenizers (Piston Pellet from Kimble Chase, Fisher Sci.) and serial dilutions were performed to determine bacterial loads.

### Statistical analysis

Data were analysed using T test non-parametric Mann-Whitney with the Graph Prism 5.0 software (San Diego, CA, USA). P-values ≤0.05 were considered significant.

