## Supplementary figure S1 for "Bread feeding is a robust and more physiological enteropathogen administration method compared to oral gavage"

### Slide 1
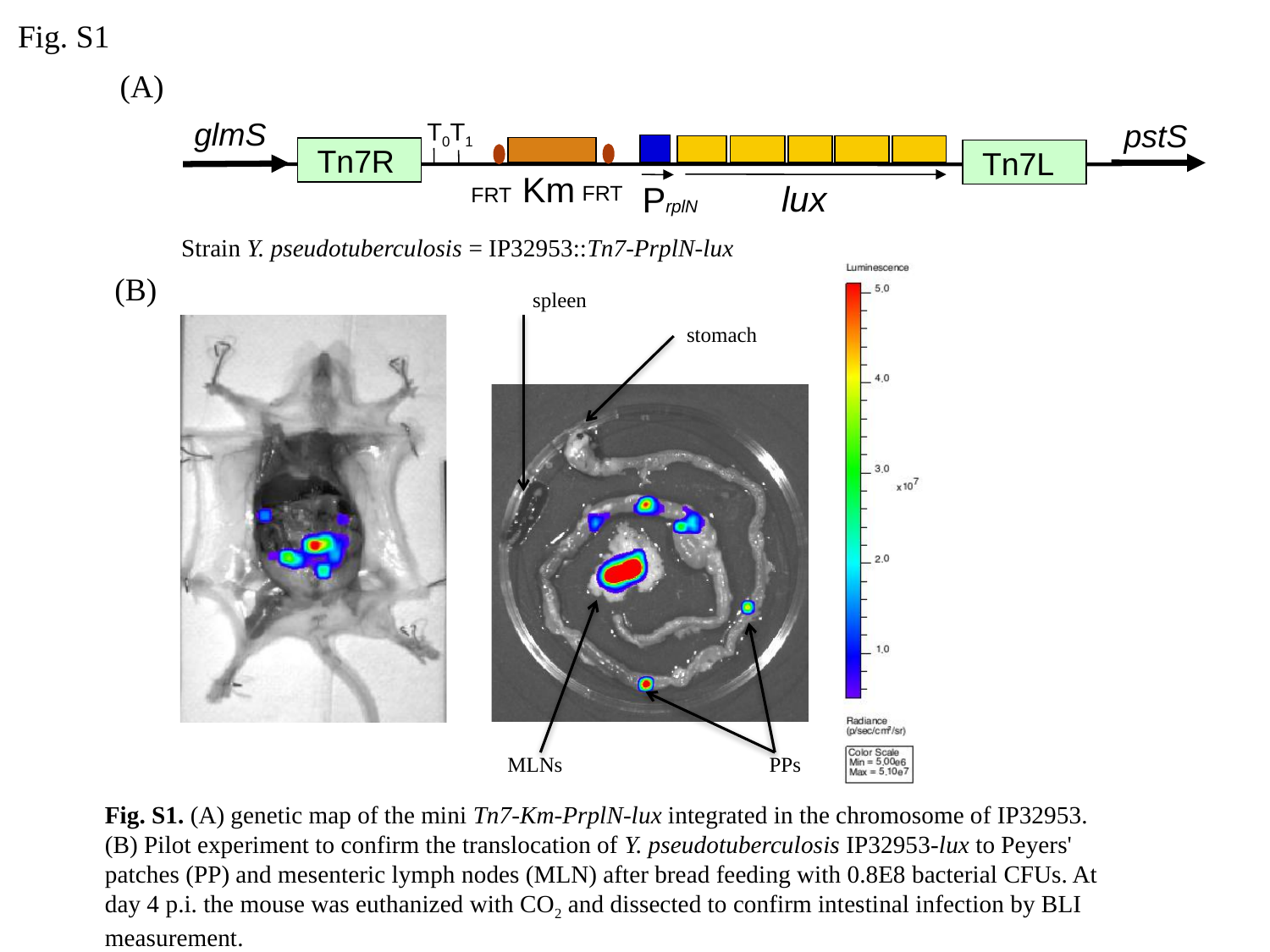

Fig. S1
(A)
glmS
pstS
T_0_T_1_
Tn7R
Tn7L
Km
lux
PrplN
FRT
FRT
Strain Y. pseudotuberculosis = IP32953::Tn7-PrplN-lux
(B)
spleen
stomach
PPs
MLNs
Fig. S1. (A) genetic map of the mini Tn7-Km-PrplN-lux integrated in the chromosome of IP32953. (B) Pilot experiment to confirm the translocation of Y. pseudotuberculosis IP32953-lux to Peyers' patches (PP) and mesenteric lymph nodes (MLN) after bread feeding with 0.8E8 bacterial CFUs. At day 4 p.i. the mouse was euthanized with CO_2_ and dissected to confirm intestinal infection by BLI measurement.
